# Time-to-onset and temporal dynamics of EEG during breath-watching meditation

**DOI:** 10.1101/2025.02.11.637771

**Authors:** Saketh Malipeddi, Arun Sasidharan, Rahul Venugopal, Prejaas K.B. Tewarie, P.N. Ravindra, Georg Northoff, Steven Laureys, Balachundhar Subramaniam, Bindu M Kutty

## Abstract

**Introduction:** Mind-body practices, such as meditation, enhance mental well-being. Research studies consistently demonstrate improved brain function and psychological well-being in meditation practitioners. A substantial body of neuroscientific evidence highlights changes in alpha and theta frequency bands during meditation among practitioners. Neurophysiological effects of meditation are reported as average power changes from resting to meditative states. However, there is a notable gap in research concerning the time-to-onset and temporal dynamics of these changes during meditation.

**Method:** Our study addresses this gap by recording high-density 128-channel EEG data during breath-watching meditation in three groups: meditation-naïve controls (n = 28), novice meditators (n = 33), and advanced meditators (n = 42). Meditators were trained in the Isha Yoga tradition. Real-time changes in brain power across different frequency bands were analyzed by segmenting the EEG data into 1-minute intervals. Using the first 30 seconds of breath-watching as the baseline, we calculated within-group power differences between this baseline and successive 1-minute segments (non-overlapping, non-sliding windows). For between-group comparisons, we assessed power differences among the three groups at 0.5, 3, 6, and 9 minutes.

**Results:** Our results indicate that time-to-onset of statistically significant increases in alpha, theta, and beta1 power, as well as decreases in delta and gamma1 power, occur around the 2-3 minute mark, with effects starting to peak between 7- and 10-minutes duration across all three groups. Statistically significant differences were observed between groups in the magnitude of these changes: advanced practitioners exhibited higher theta and theta-alpha power at all time points compared to the other groups.

**Conclusion:** Our findings suggest that neurophysiological changes begin around 2-3 minutes after starting meditation and peak around 7-10 minutes across all three groups. However, the magnitude of these effects is greater in the advanced meditator group. As long as meditation retreats are not possible for many individuals, brief meditation practices of 7 minutes or more, delivered through digital platforms, could offer accessible, effective, and scalable solutions to improve mental well-being. This suggests a broader application of meditation practices in daily life, encouraging even those with tight schedules to incorporate such beneficial practices.

## Introduction

Meditation has been the most popular mind-body intervention (an increase from 7.5 to 17.3 percent in the US from 2002 to 2022 and practiced by 200-500 million people globally) ^1–3^, practiced for improving attention, awareness, and enhancing emotion regulation ^4–6^.

Importantly, meditation has been shown to lower psychological stress, anxiety, and depression and enhance joy, compassion, resilience, and well-being ^7–13^.

Meditation includes a wide variety of practices, popularly classified into focused-attention, open-monitoring, and loving-kindness practices ^14–16^. Focused-attention practices include those practices that train attention on a particular object, such as the breath ^16–18^. This simple aspect of paying attention to the breath, called breath watching or mindful breathing, has been used in Buddhist and Yogic traditions to overcome suffering and gain insight into the true nature of the self ^11,19,20^. In addition to training attention, these practices also train awareness. Whenever the mind wanders, individuals need to cultivate awareness to notice mind-wandering and bring the attention back to the breath. Attention and awareness training, which are impaired in many mental health disorders, have been shown to be crucial for well-being ^4–6,16,17,21^.

The meditators practiced Isha Yoga, an international school of Yoga that provides tools for enhancing overall well-being. Research on Isha Yoga practices shows beneficial outcomes, including reduced stress levels, heightened mindfulness, and enhanced well-being ^10,13,22–24^. These practices have also demonstrated benefits such as better cardiac sympathovagal balance ^25^, heightened visual attention ^26^, elevated brain-derived neurotrophic factor (BDNF) and cortisol awakening response ^27^, increased anandamide levels ^28^, and more ^29,30^. What are the neurodynamic features mediating the effects of Isha Yoga? While some studies used fMRI^31^ and EEG^32,33^, the dynamic neural changes during the actual meditative state itself remain yet unclear. Addressing this gap in our knowledge is the goal of our study.

Electroencephalography (EEG), with its excellent temporal resolution, has been extensively used to study meditation ^14,30,32,34,35^. Research consistently shows an increase in power in alpha and theta frequency bands in the resting state (as a trait effect) and during meditation (as a state effect) ^14,35^. Changes in other bands, such as gamma, are reported only among advanced meditation practitioners ^33,36,37^. Further, meditators experience a state of ‘relaxed alertness’ during meditation, where the mind is relaxed and alert at the same time. This state is reflected in the EEG by an increase in power in both low-frequency and high-frequency bands ^32,35,38,39^.

Research so far has treated the brain during meditation as a static entity, while ignoring the potential dynamic modulations of brain activity during (the initiation of) the meditative state. It is common to analyze EEG data by examining short epochs, which are then averaged to obtain average activity^14,35^. For instance, in a study on Vipassana meditation (40 mins), the authors took mid-2-minute EEG epochs every 10 mins of the data and then averaged the power values ^40^. In one of our prior studies, we analyzed EEG data from breath-watching meditation by selecting 2-second epochs and averaging them to discern overall power differences (rest to meditation, between different groups) ^32^. These approaches effectively reduce the computational demands associated with analyzing extensive EEG data. However, they may inadvertently neglect the critical aspects of the time-to-onset of neurophysiological effects and the intricate temporal dynamics of brain oscillation changes that occur throughout the meditation practice period. This oversight could limit our understanding of how these neural patterns evolve over time in response to meditation. This information may better help us to customize the duration of meditation practice for beginners.

Unlike these approaches, we here aim to investigate the dynamic neural changes during a meditation state itself: how does neural brain activity like the power of its different oscillations (in delta, theta, alpha, beta and gamma) change during the course of a meditative state? Specifically, using EEG in a dynamic way, we sought to answer three key questions as aims: 1) When do modulations in brain activity appear after starting meditation? 2) When do these modulations peak? and 3) Are there differences in these oscillatory changes between meditation-naïve controls, novice meditators, and advanced practitioners?

## Methods

### Experimental setup

We performed the analyses on the data already published by us ^32^. EEG data were recorded using a 128-channel HydroCel Geodesic Sensor Net from the Geodesic EEG System 300 (Philips Neuro, USA) with a sampling rate of 1 kHz and digitized at 24-bit resolution. The procedure was conducted in a soundproof room with an ambient temperature of 25°C and humidity controlled between 40% and 60%. Participants were instructed to wash their hair before the session to ensure high-quality EEG signals. Electrode impedance was maintained below 50 kΩ, as recommended by the vendor. For more details about the dataset, such as data collection and procedure, the reader is directed to ^32^.

### Participants

Individuals with no previous meditation practice (meditation-naïve controls) were recruited (*n* = 28, 16 females, mean age (SD) = 31.14 (6.38)) from the local community. Instructions related to breath-watching meditation were given before the session. Meditators were recruited from the Isha Yoga centers in Bangalore, Karnataka. They were classified as novice (n = 33, 14 females, mean age (SD) = 31.66 (7.64)) or advanced meditators (n = 42, 18 females, mean age (SD) = 35.57 (6.81)) based on completion of Samyama, an advanced meditation retreat. Inclusion criteria for all participants included: age range 25-50 years; healthy or under stable dosage of non-psychoactive medication (stable dosage period considered as a minimum of one month with no changes anticipated); able to read, write, and speak fluently in English, and able to comprehend and answer standard questionnaires.

Exclusion criteria for all participants included a history of neurological disease, vision defects (uncorrected), auditory deficits or severe physical disabilities, history of substance abuse/dependence, history of major mental illness, any psychiatric medication, or psychotherapy. Novice and advanced meditators had to meet the following exclusion criteria: no current practice or history of practice of any other school of Yoga or meditation. The estimated lifetime hours of meditation practice of the groups are given in Table 1. This study received approval from the Institutional Human Research Ethics Committee (NIMH/DO/ETHICS SUB-COMMITTEE MEETING/2018). All participants provided written informed consent before participating in the study, and no monetary compensation was offered.

**Table 1.** Estimated lifetime hours of Isha Yoga practice.

| Lifetime hours of Isha Yoga practice |  |  |  |
| --- | --- | --- | --- |
|  | Controls | Novice meditators | Advanced meditators |
| Hours (mean) | 0 | 1637 | 5508 |
| Hours (std. deviation) | 0 | 1127 | 2897 |

### Procedure

The entire experimental paradigm consisted of multiple sessions in the following sequence: resting state (4 minutes), sukha kriya pranayama (6 minutes), breath-watching meditation (15 minutes), resting state (4 minutes), visual oddball task (15 minutes), resting state (4 minutes), shoonya meditation (15 minutes), and resting state (4 minutes). These details are described in our previous research ^32^. In this study, we focused on investigating ongoing temporal dynamics of oscillations during breath-watching meditation. During this session, participants paid attention to the natural movement of their breath without identifying with their thoughts. Meditators performed breath-watching as they typically do in their regular practice, while meditation-naïve controls (who are first-time meditators) were taught this practice before the session (Practice is freely available online: https://www.youtube.com/watch?v=C_xsXnRd_uc).

### Pre-processing

EEG data preprocessing was performed using the open-source MATLAB R2021b toolbox EEGLAB v2021.0 ^41^. The EEG data were resampled from 1024 Hz to 250 Hz using EEGLAB’s resample function and re-referenced to the average reference. The data were then band-pass filtered between 0.5 Hz and 40 Hz. Subsequently, bad channels and segments were automatically identified and rejected using the Artifact Subspace Reconstruction (ASR) plugin, with a threshold of five standard deviations. Finally, stationary artifacts, including eye movements, breathing artifacts, and muscular noise, were reduced using the ICALABEL plugin (90% threshold) after running independent component analysis (ICA) with the ’infomax’ method on the cleaned data.

### EEG analyses

Power spectral values (using the ‘bandpower’ function; 0.5 Hz resolution; ‘Hamming’ window) were extracted for standard frequency bands: delta (1–4 Hz), theta (4–8 Hz), theta-alpha (6-10 Hz), alpha (8–12 Hz), beta1 (13–20 Hz), beta2 (20–30 Hz), and gamma1 (30–40 Hz) during breath-watching meditation. We used the first 30 seconds of breath-watching as the baseline to explore the time-to-onset of neurophysiological effects and the temporal dynamics of the practice across all three groups. Time-to-onset was defined as the time point at which significant brain changes occur. For within-group comparisons, differences in absolute power between the first 30 seconds of data and successive 1-minute segments (non-overlapping non-sliding windows) were calculated. Each 1-minute time point represented cumulative power until that point. For between-group comparisons, differences in absolute power among the three groups at 0.5 minutes, 3 minutes, 6 minutes, and 9 minutes were calculated.

### Statistical analyses

Statistical comparisons were conducted using the LIMO toolbox ^42^, employing robust t-statistics with 2000 permutations, with corrections for multiple comparisons across electrodes using cluster statistics. Statistical significance was set at alpha < 0.05.

## Results

### Within-group comparisons of dynamic changes over time during mediation practice

Within the meditation-naïve controls, novices, and advanced meditators, time-to-onset of neurophysiological effects was observed around 2-3 minutes (Table 2). Compared to the first 30 seconds of breath-watching meditation, significantly increased power in theta, alpha, and beta1 bands and significantly decreased power in delta and gamma1 bands were found in all three groups. No changes were detected in the beta2 band in any group. Differences in the time-to-onset of effects were seen across frequency bands and between groups.

**Table 2:** Temporal and Power Dynamics of EEG Effects Across Frequency Bands and Groups during Breath-Watching meditation.

| Frequency band | Group | Time-to-Onset (min) | Maximum effects (min) | Power change from first 30 seconds (Increase/Decrease) ($p < 0.05$) |
| --- | --- | --- | --- | --- |
| **Delta (0.5 – 4 Hz)** | Controls | 2 | 10 | Decrease |
|  | Novice meditators | 1 | 10 | Decrease |
|  | Advanced meditators | 2 | 9 | Decrease |
| **Theta (4 – 8 Hz)** | Controls | 2 | 8 | Increase |
|  | Novice meditators | 2 | 7 | Increase |
|  | Advanced meditators | 1 | 10 | Increase |
| **Theta-alpha (6 – 10 Hz)** | Controls | 2 | 10 | Increase |
|  | Novice meditators | 2 | 10 | Increase |
|  | Advanced meditators | 2 | 10 | Increase |
| **Alpha (8 – 12 Hz)** | Controls | 2 | 10 | Increase |
|  | Novice meditators | 2 | 10 | Increase |
|  | Advanced meditators | 2 | 9 | Increase |
| **Beta1 (13 – 30 Hz)** | Controls | 2 | 7 | Increase |
|  | Novice meditators | 1 | 9 | Increase |
|  | Advanced meditators | 2 | 8 | Increase |
| **Gamma1 (30 – 40 Hz)** | Controls | 2 | 10 | Decrease |
|  | Novice meditators | 3 | 10 | Decrease |
|  | Advanced meditators | 3 | 10 | Decrease |

For the control group (Figure 1), significant time-to-onset of effects was observed at 2 minutes in the delta band (central, left parieto-occipital, and right fronto-parietal regions), 2 minutes in the theta band (left parieto-occipital and right fronto-parietal regions), 2 minutes in the theta-alpha band (whole brain), 2 minutes in the alpha band (whole brain except right central), and at 2 minutes in the beta1 and gamma1 band (whole brain).

**Figure 1.**
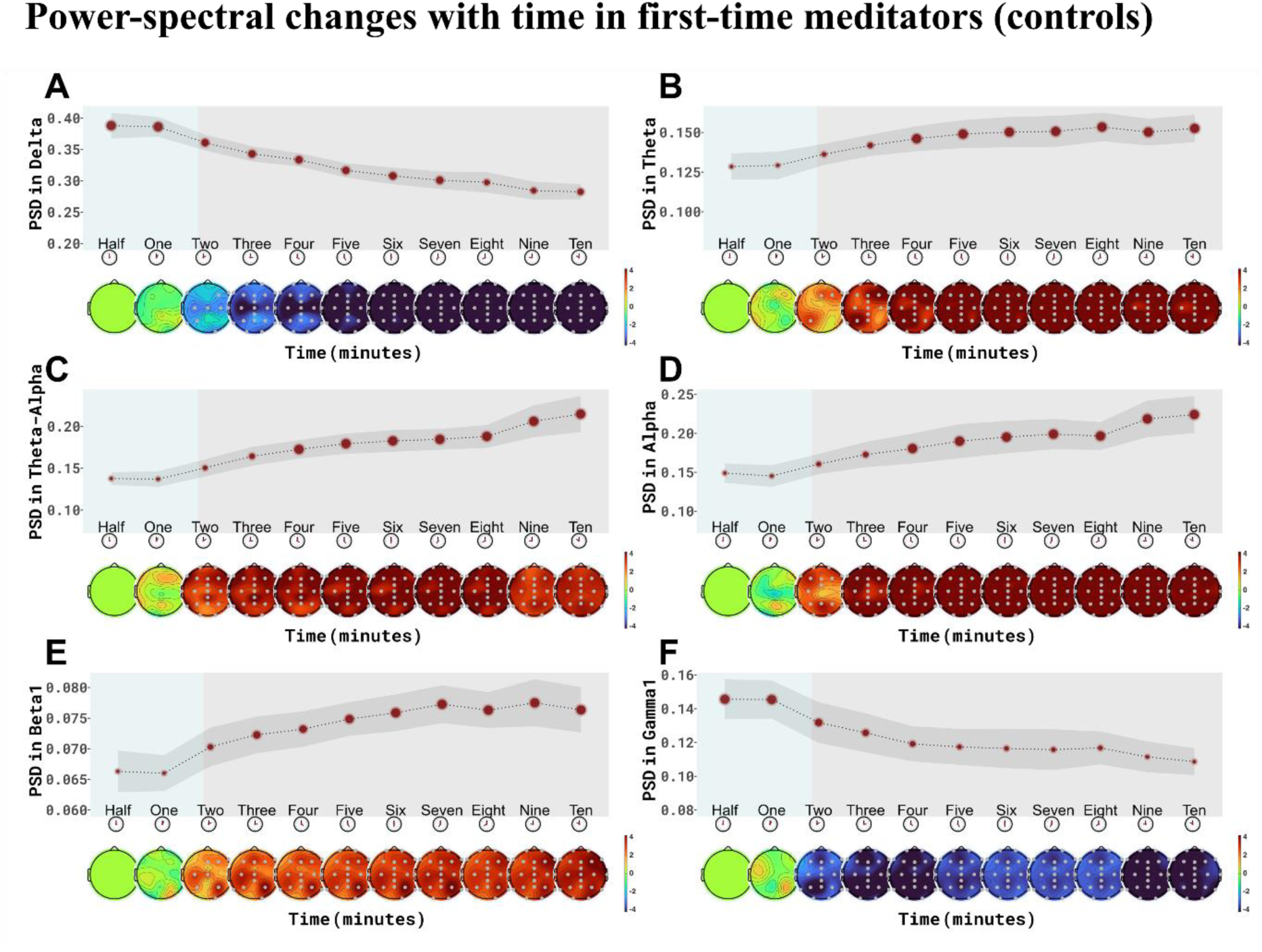
Time to onset in meditation-naïve controls: Time-to-onset and temporal dynamics during breath-watching meditation in first-time meditators (meditation-naïve controls) (CNT) are illustrated. Power-spectral values in different frequency bands show time-to-onset of effects around a 2-3-minute duration. The size of the dots corresponds to the magnitude of the values. The grey bar indicates standard error. The lower head plots are the t-value Topo-plots (compared to the first 30 seconds) that highlight electrodes with statistically significant differences (dots, p<0.05) in power. P-values have been corrected for multiple comparisons using cluster-based statistics.

In the novice meditator group (Figure 2), different time-to-onset of effects was observed. The electrodes showing significant differences were also different compared to controls.

**Figure 2.**
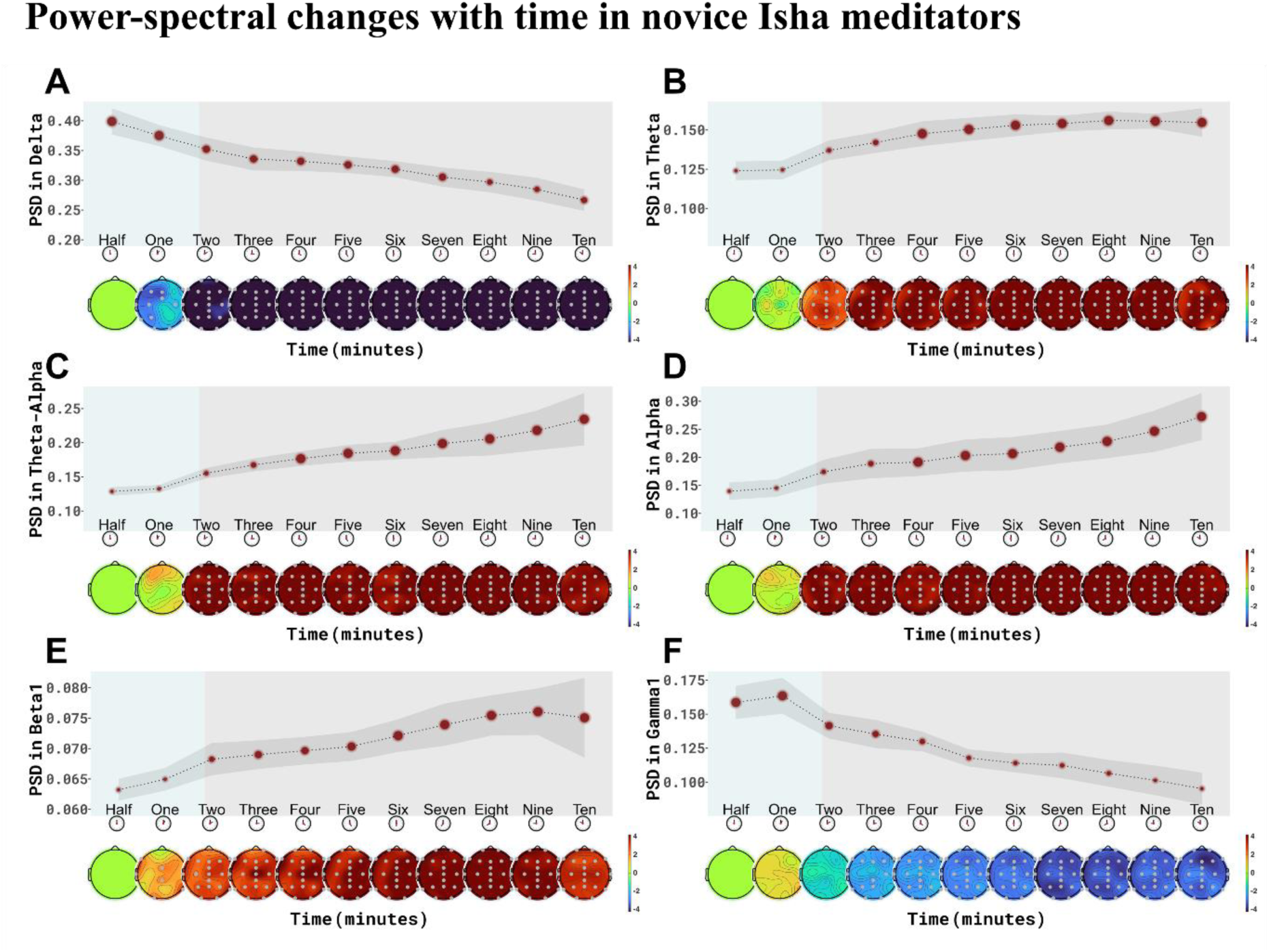
Time to onset in novice meditators: Time-to-onset and temporal dynamics during breath- watching meditation in novice meditators (NOV) are illustrated. Power-spectral values in different frequency bands show time-to-onset of effects around a 3-minute duration. The size of the dots corresponds to the magnitude of the values. The grey bar indicates standard error. The lower head plots are the t-value Topo-plots (compared to the first 30 seconds) that highlight electrodes with statistically significant differences (dots, p<0.05) in power. P-values have been corrected for multiple comparisons using cluster-based statistics.

Significant time-to-onset of effects was observed at 1 minute in the delta band (whole brain except right posterior regions), 2 minutes in the theta, theta-alpha, and alpha bands (whole brain), 1 minute in the beta1 band (fronto-central and central regions), and 3 minutes in the gamma1 band (whole brain).

Finally, in the advanced meditator group (Figure 3), significant time-to-onset of effects was noticed at 2 minutes in the delta band (whole brain), 1 minute in the theta band (whole brain), 2 minutes in the theta-alpha band (posterior regions), 2 minutes in the alpha band (whole brain), 2 minutes in the beta1 band (posterior regions), and 3 minutes in the gamma1 band (whole brain).

**Figure 3.**
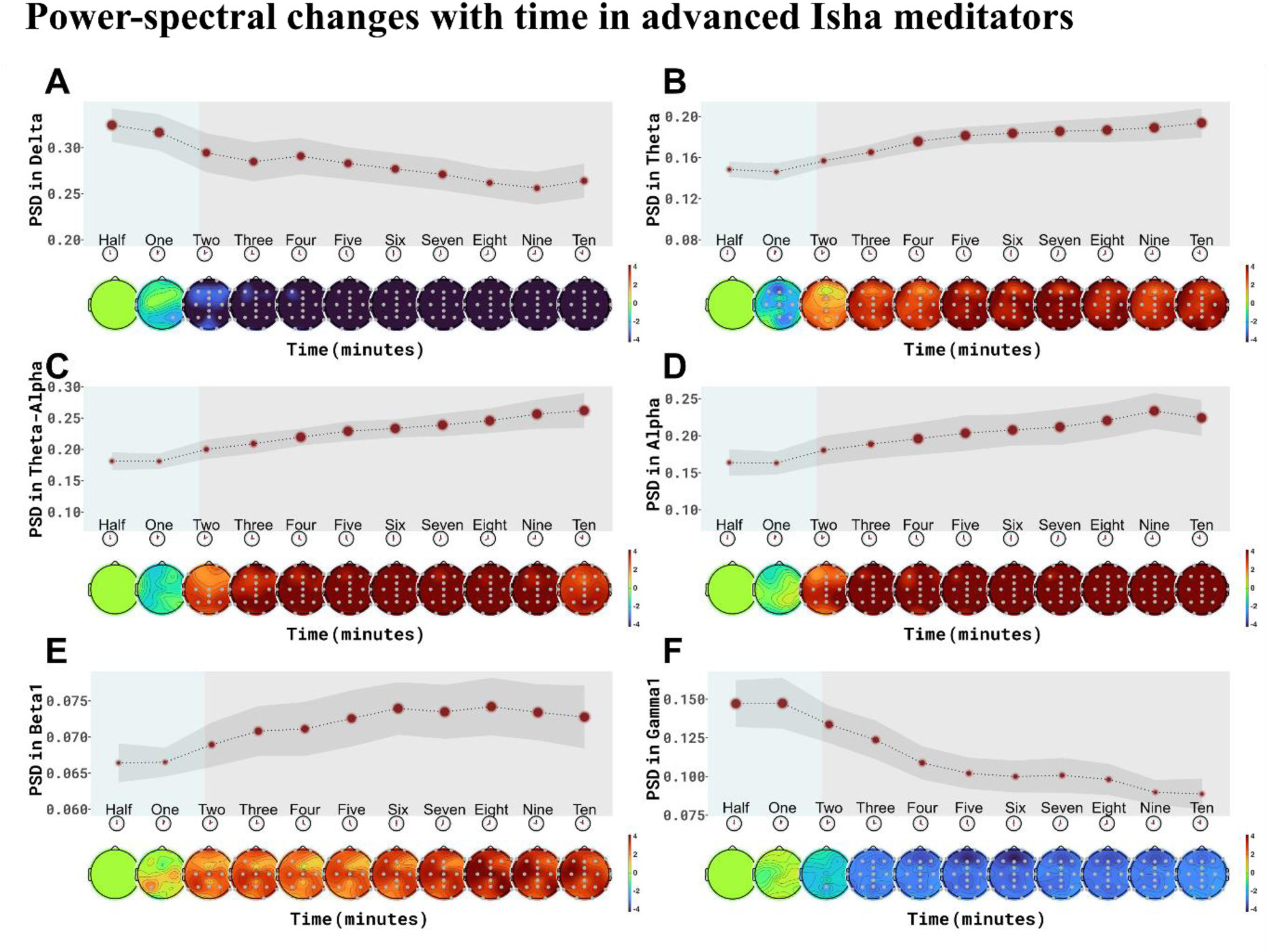
Time to onset in advanced meditators: Time-to-onset and temporal dynamics during breath-watching meditation in advanced meditators (ADV) are illustrated. Power-spectral values in different frequency bands show time-to-onset of effects around a 2-minute duration. The size of the dots corresponds to the magnitude of the values. The grey bar indicates standard error. The lower head plots are the t-value Topo-plots (compared to the first 30 seconds) that highlight electrodes with statistically significant differences (dots, p<0.05) in power. P-values have been corrected for multiple comparisons using cluster-based statistics.

For all the three groups, delta and gamma1 power decreased with time, whereas theta, theta-alpha, alpha, and beta1 power increased with time. The maximal effects in power were observed in the 7–10-minute duration in the different groups. These results are summarized in Table 2.

In summary, all groups showed dynamic changes in the power of different frequency including delta (decrease), theta and alpha (increase) and gamma (decrease) which started around 2-3 min and peaked around 7 mins. The exact timing, power bands, and electrodes differed between the three groups, though, showing either earlier or later changes. We therefore conducted between-group comparisons in the next step.

### Between-group comparisons

Our results showed significantly lower power in the delta band in advanced meditators at 0.5 minutes compared to novice meditators and at 0.5- and 3 minutes compared to controls (Figure 4 (A)). Additionally, advanced meditators showed significantly higher theta power than both groups at the 0.5-, 3-, 6-, and 9-minute durations (Figure 4 (B)). Similarly, advanced meditators showed significantly higher power in the theta-alpha band at 0.5-, 3-, and 6-minute durations compared to controls, and at 0.5 minutes compared to novice meditators (Figure 4 (C)). In the gamma frequency band, advanced meditators showed lower power than controls at 9 minutes, with no significant differences observed at other time points (Figure 4(D)). There were no statistically significant differences in other bands or comparisons between novices and controls. Overall, our results indicate that while advanced meditators do not exhibit differences in time-to-onset or peak EEG effects compared to other groups, the magnitude of these effects is greater. Specifically, advanced meditators showed stronger decreases in delta and gamma as well as stronger increases in alpha and theta compared to the other two groups.

**Figure 4.**
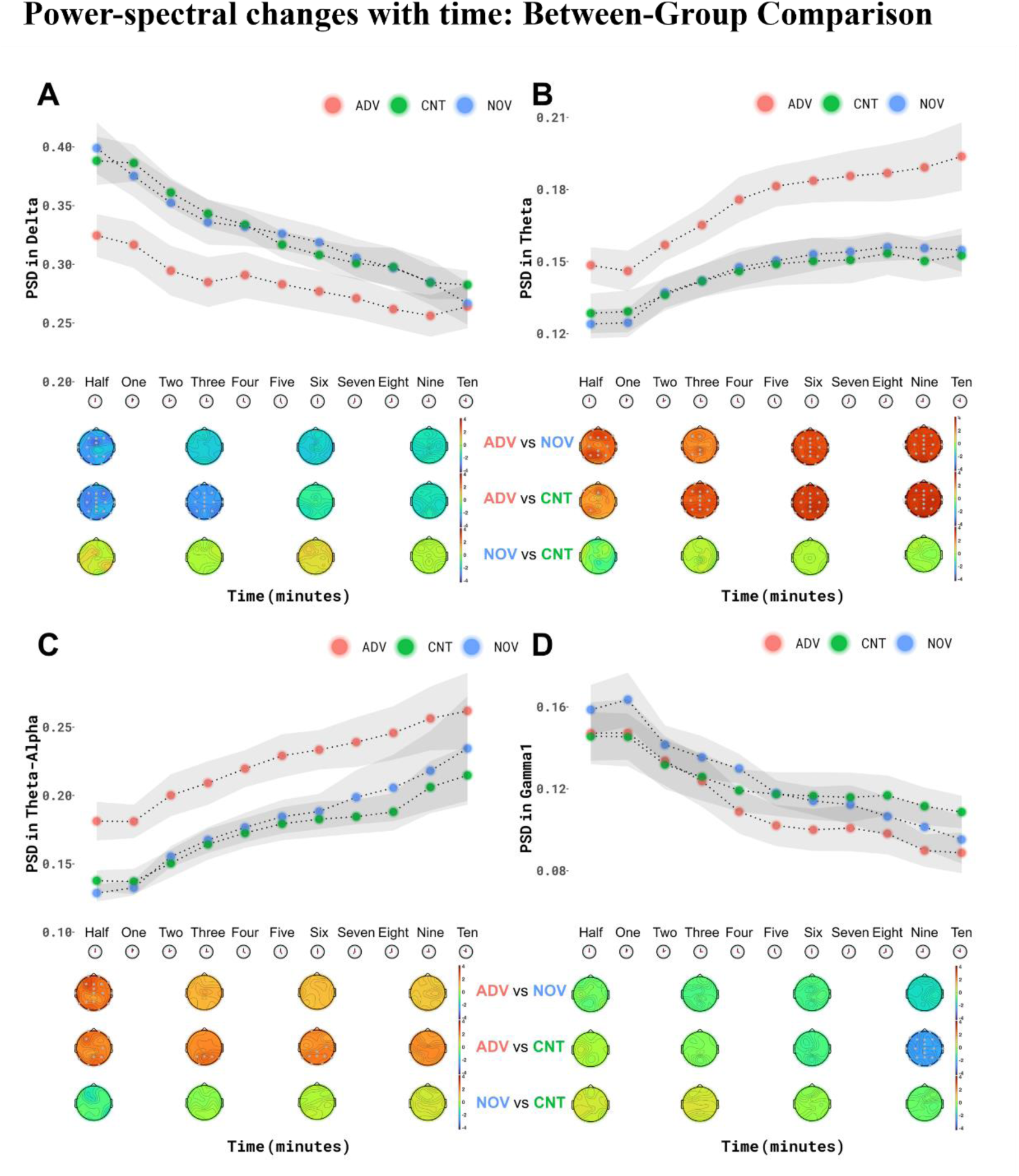
Between group comparisons: Between-group comparisons of the temporal dynamics of neurophysiological effects across delta (A), theta (B), theta-alpha (C), and gamma (D) frequency bands during breath-watching meditation at 0.5, 3, 6, and 9 minutes. The grey bar shows the standard error of the mean. The lower head plots are the t-value Topo-plots that highlight electrodes with statistically significant differences (dots, p<0.05) in power. P-values have been corrected for multiple comparisons using cluster-based statistics. Legend: ADV = Advanced meditators, NOV = Novice meditators, CNT = Meditation-naïve controls, PSD = Power-spectral density.

Overall, our results show: (1) decrease in delta and gamma over the course of 10 min meditation practice in all three groups starting around 2 min and peaking around 7-10 min; increase in alpha and theta power the course of the 10 min meditation practice starting around 2 min and peaking around 7-10 min in all three groups; and (3) significantly different power values in both delta/gamma (lower) and alpha/theta (higher) in the advanced mediators compared to the other two groups.

## Discussion

Our study used a novel approach by examining the dynamic changes in the oscillatory power in EEG during breath-watching meditation in advanced meditators, novice meditators, and controls. We sought to answer three key questions: (1) When do modulations in brain activity appear after the onset of meditation? We showed a time-to-onset of spectral effects (an increase in alpha, theta, and beta1, and a decrease of delta and gamma1 power) following breath-watching meditation around a 2–3-minute duration in all three groups. (2) When do these modulations peak? Effects peaked around the 7–10-minute duration in all the groups. (2) Are there differences in these oscillatory changes between meditation-naïve controls, novice meditators, and advanced practitioners? Our findings revealed unique differences between the groups in these temporal effects. Advanced meditators showed higher theta and theta-alpha power and lower delta and gamma power than the other groups.

Our first main finding shows an increase in theta, alpha, and beta1 power and a decrease in delta and gamma1 power in all three groups at a 2–3-minute duration. In other words, this suggests that significant neurophysiological changes may take a couple of minutes to appear after starting meditation. This could mean that the mind takes some time to settle down into a meditative state, even for advanced meditators. Enhanced theta and alpha power are indicators of mental relaxation, a widely reported feature of many different meditation practices ^14,35,43–45^. Greater beta1 power could indicate alertness ^35,39,46^. Overall, a simultaneous increase in low-frequency alpha and theta and high-frequency beta1 power could mean a state of relaxed alertness in the practitioners ^32,35,39^. A decrease in delta and gamma1 power with time is more difficult to interpret as there are mixed findings in the literature ^35^, but could mean more alertness ^47^ and lower mind-wandering ^48^, respectively.

Alongside observing significant changes around the 2–3-minute mark, our findings show peaking of neurophysiological effects around the 7–10-minute duration. Information like this cannot be gleaned from static perspectives. It is important to interpret our findings in the context of the baseline state used. While the studies mentioned earlier used the eyes-closed resting state as their baseline, we employed the first 30 seconds of meditation practice as our baseline. This approach is justified, as our objective was to examine temporal effects, which required maintaining the same state as the baseline for consistency and relevance in the comparisons.

Given this information, how does expertise in meditation relate to these changes? To this end, we looked at the differences between the practitioners. Although no major differences were seen between the groups in the onset and peaking of neurophysiological effects, the magnitude of changes differed.

Advanced meditators showed significantly greater theta power at all timepoints, with higher theta power observed as early as 30 seconds. This indicates a trait feature of meditation ^32^. Theta power increased linearly until 6 minutes before plateauing, whereas in the other two groups, plateauing occurred between 3 and 6 minutes. This could mean that advanced meditators achieved deeper states of meditation as time progressed. Theta power is a marker of expertise in meditation, with several studies reporting greater theta power in advanced meditators, compared to novice meditators, during meditation ^14,34,35,43^. Higher power is linked to greater synchronization of neural oscillations ^16,43,49^. Given that breath-watching is a focused-attention practice, and advanced meditators are more trained in their attentional skills, attentional networks may be more synchronized, leading to greater theta power. In fact, in our previous research based on the same data, we showed that advanced meditators reported fewer hindrances (such as mind-wandering), higher levels of concentration (such as more focused attention), and an overall greater depth of meditation compared to the other groups ^32^. Overall, higher theta power in advanced meditators could indicate greater attention, meta-awareness, and pleasant states like bliss ^35,44,50,51^.

Lower delta power in advanced meditators compared to the other groups could indicate a more alert and aware state of meditation ^34,47^. Lower gamma power observed in advanced meditators compared to controls at 9 minutes may indicate that, as time progresses, meditation deepens, leading to less mind-wandering and greater focus ^48^. Finally, a large body of research documenting brain changes in short-term (like our novice practitioners) and long-term meditators (like our advanced practitioners) from various traditions ^14,34,35^. However, limited research has explored brain changes in first-time meditators (like our controls) ^52^. Our findings show that meditator-like neurophysiological changes are interesting and worth exploring in future research.

Participation in long meditation retreats has been shown to significantly enhance well-being ^11,27,28^. However, this may not be practical or feasible for large segments of the population. Digital interventions that provide accessible meditation training through apps or online programs could help bridge this gap, making meditation more accessible to a wider population. In this context, our findings that just 3–7 minutes of meditation can induce significant neurophysiological changes, even in first-time meditators like our controls, are encouraging. Such brief meditation sessions could maximize benefits while minimizing common challenges faced by beginners, such as cognitive fatigue, boredom, and drowsiness. These findings support the integration of brief meditation practices into daily routines for mental health and cognitive benefits.

Mental health issues such as stress, anxiety, and depression are rising at an alarming rate worldwide ^53–56^. Research indicates that one in two individuals will develop a mental health disorder by the age of 75 ^57^. In response to this growing crisis, the World Health Organization’s Comprehensive Mental Health Action Plan 2013-2030 emphasizes the importance of investing in prevention and promoting mental well-being ^58^. Given growing scientific evidence supporting the role of meditation in promoting mental well-being ^7,8,10,12,23,24,59^, and the recent declaration of December 21st as World Meditation Day by the United Nations General Assembly ^3^, which underscores the relevance and importance of meditation, our findings advocate for the global implementation and widespread adoption of simple meditative practices to nurture a healthy mind.

## Limitations

An important limitation of this study is the lack of phenomenological data on temporal changes in meditative depth and quality, which could have complemented the neurophysiological findings and offered deeper insights into the temporal dynamics of meditation. Additionally, the absence of pre- and post-meditation mental well-being measures limited us from correlating frequency band changes with subjective reports of mental well-being.

## Conclusion

Our study addresses a critical gap in the literature by providing insights into when neurophysiological changes begin and peak during meditation. We observed that these changes start around 2–3 minutes and peak around 7-10 minutes across all groups during breath-watching meditation. These changes included increased alpha, theta, and beta1 power and decreased delta and gamma1 power. Notably, while meditation expertise did not alter the timing of these changes, it significantly influenced their magnitude, with advanced meditators showing greater theta power and lower delta and gamma power. Overall, we suggest that the brain’s response to meditation can be rapid and varies with the practitioner’s experience, potentially influencing cognitive and emotional processing in significant ways. Our findings offer novel insights into the temporal dynamics of EEG changes during meditation. The study underscores the potential of personalized meditation programs tailored to individual experience levels, optimizing mental health outcomes.

## Ethics Statement

All procedures performed in studies involving human participants were per the ethical standards of the institutional research committee and with the 1964 Helsinki Declaration and its later amendments or comparable ethical standards. The study was approved by the NIMHANS Human Ethics Committee (NIMH/DO/ETHICS SUB-COMMITTEE MEETING/2018).

## Informed Consent statement

Freely given, written informed consent to participate in the study was obtained from participants.

## Acknowledgments

We express our gratitude to Maa Vama, Isha Foundation, for the support and guidance in conducting this research and facilitating Isha meditators recruitment.

